# CODABEILLES: A Reliable Reference Library of COI DNA Barcodes for French Wild Bees Monitoring (Apoidea: Anthophila)

**DOI:** 10.1101/2025.11.10.687545

**Authors:** M. Ollivier, A. Marquisseau, E. Dufrêne, R. Rudelle, The CODABEILLES Consortium, R. Rougerie, A. Perrard, M. Pichon

## Abstract

In the Anthropocene, the decline of insect pollinators poses a significant threat to ecosystem services, particularly to wild bee populations essential for plant biodiversity and agricultural productivity. France, with 983 species, hosts one of the most diverse bee faunas in Europe, yet these species face growing pressures from habitat loss, climate change, and intensive agriculture. Addressing this crisis requires robust taxonomic frameworks and efficient species identification methods to support long-term monitoring initiatives such as the European Pollinator Monitoring Scheme, EU-PoMS.

DNA barcoding, utilizing the COI-5P gene, has proven effective for species delineation and biodiversity monitoring, particularly in detecting cryptic diversity among genera with large numbers of species such as *Andrena, Nomada* or *Lasioglossum*. However, significant gaps remain in reference libraries, particularly for the species from the Mediterranean Basin. To bridge this gap, the CODABEILLES initiative was launched in 2021 to enhance barcode data for the French bee fauna. Initially, only 25% of species had barcodes from French voucher specimens, increasing to 62% when considering voucher specimens from other countries. By 2025, thanks to collaboration with sixteen specialists and institutions, CODABEILLES contributed 1477 reference barcodes, covering approximately 560 species and raising barcode coverage to 82%. When integrating data published under other initiatives over the same period the coverage reaches 94% of the French bee fauna. This dataset significantly enhances species identification accuracy and supports large-scale pollinator monitoring through metabarcoding and environmental DNA approaches.

Despite the success of COI-5P barcoding, taxonomic inconsistencies persist, necessitating further integrative research. This study underscores the need for continued collaboration among taxonomists, molecular biologists, and conservationists to refine species classifications and ensure comprehensive reference databases. The improved barcode coverage provided by CODABEILLES paves the way for more accurate DNA-based monitoring of wild bee populations and their ecological interactions, crucial for guiding conservation strategies in the face of ongoing environmental change.

## Background

In the Anthropocene, growing evidence of massive declines in insect populations (in abundance, biomass, diversity, or spatial distribution) poses a threat to ecosystem services (Wagner et al. 2021). Animal pollination, mainly provided by a diversity of wild bee species (Hymenoptera, Apoidea, Anthophila), is essential for the production of 75% of cultivated plants but also for the conservation of wild plant biodiversity (Klein et al. 2007; Potts et al. 2016; Goulson 2019). Wild bees are a diverse group among hymenopterans, counting over 20,000 species worldwide (Ascher and Pickering 2025). Global bee distribution shows a specific pattern: while tropical environments are renowned for their extraordinary richness in many insect species, relatively xeric areas surpass other regions in terms of bee species richness (Orr et al. 2021; Leclercq et al. 2023). The Mediterranean Basin is therefore considered a hotspot of bee diversity (Wood et al. 2024; Schneider et al. 2024). With 983 species, France (Mainland and Corsica) harbours the 4^th^ most diversified fauna in Europe after Greece, Spain and Italy (Reverté et al. 2023; Ropars et al. 2025). This accounts for nearly half of Europe’s 2,138 bee species (Reverté et al. 2023).

The loss of insect pollinators is attributed to a combination of global factors that cause significant disruptions to wildlife. Among the well-documented drivers of pollinator collapse are climate change and land use change, which result in the loss of natural habitats. This trend affects both bumblebees (White and Dillon 2023; Singh et al. 2024; Ghisbain et al. 2024) and solitary bees (LeBuhn and Vargas Luna 2021) as today’s agricultural landscapes fail to provide adequate nesting sites and floral resources (Goulson et al. 2015; Carrié et al. 2018). Intensive agriculture, often linked to the use of pesticides and fertilizers, is another significant stressor contributing to the decline of wild bees (Goulson et al. 2015). However, the local impacts of agricultural practices can be mitigated at landscape scale by the presence of semi-natural habitats such as woods and grasslands (Park et al. 2015; Carrié et al. 2017; Rivers-Moore et al. 2020). The incorporation or restoration of flower-rich areas into farmland is a practical measure that should be enhanced. For instance, the research project RestPoll (Restore Pollinator Habitats in Europe), along with agri-environmental schemes at both national and European levels (French “Plan national en faveur des insectes pollinisateurs et de la pollinisation 2021-2026” and European Pollinator Monitoring Scheme, EU-PoMS, JRC138660), represent recent initiatives aimed at addressing a shared challenge: counteracting or slowing the collapse of pollinator populations in the coming decades.

The Europe-wide long-term monitoring conducted under EU-PoMS relies on standardized sampling methods (Potts et al. 2020). Like any monitoring program in insect ecology and conservation studies, it requires two critical elements to ensure effectiveness: 1) a robust taxonomic framework, and 2) an accurate, rapid and cost-effective identification of bee specimens at the species level. Despite significant efforts to advance knowledge of the European bee fauna (Ghisbain et al. 2023), new species are still frequently described (Wood 2022; Le Divelec 2024) and species boundaries within certain groups remain unresolved. DNA barcoding, which uses the 5’ region of the mitochondrial gene cytochrome c oxidase 1 (COI- 5P) as a standardized marker for species delineation, has progressively been adopted as part of an integrative taxonomy approach (Hebert et al. 2003; Miralles et al. 2024). It helped in clarifying groups of species being historically controversial and has frequently demonstrated its capacity to uncover cryptic bee species (Williams et al. 2012; Pauly et al. 2015). For instance, European bees of the genus *Andrena* subgenus *Taeniandrena* have been investigated, combining DNA barcodes and morphology, revealing an unexpected level of cryptic diversity in a geographically restricted area (Praz et al. 2022).

For several bee genera, an accurate species-level identification requires advanced taxonomic expertise (Rondeau et al. 2023). Although training opportunities are beginning to emerge, these skills remain confined to a few experts and are even unavailable in many countries. The integration of DNA barcoding with high-throughput sequencing (HTS) technologies presents a cost- and time-efficient solution for scaling up routine pollinator monitoring on a large scale (Chua et al. 2023). This approach is referred to as *metabarcoding* or *megabarcoding*, depending on how individuals are pooled during the process and the resulting ability to obtain abundance data (Taberlet et al. 2012; Gueuning et al. 2019). The accuracy of a specimen assignment to a species through DNA barcoding heavily depends on the completeness of the DNA barcode reference libraries and on the sufficient characterization of intraspecific variability (Phillips et al. 2019).

In the past decades, only a few national and regional libraries have been established for wild bees, for example in Chile (Packer and Ruz 2017), Canada (Sheffield et al. 2017), Ireland (Magnacca and Brown 2012), United Kingdom (Creedy et al. 2020), Luxembourg (Herrera-Mesías et al. 2022) and more recently Slovenia (Janko et al. 2024). Despite global efforts to develop DNA barcode references for wild bee species, the reference library for French species remains incomplete. Significant gaps exist for species from the Mediterranean Basin, barcodes from French vouchers are necessary for a better characterization of intraspecific diversity, and, where barcodes are available, often a very limited number of specimens has been sequenced. In early 2021, when the CODABEILLES initiative was launched to enhance reference data on the French bee fauna, only 25% of species were covered by barcodes obtained from French specimens, mainly collected in the Loire Valley (Villalta et al. 2021). Coverage increased to 62% when including German bee specimens provided by an extensive work by Schmidt et al. (2015). Very recently, Wood et al. (2024) made a considerable contribution to documenting wild bees in the Iberian Peninsula providing useful data for some French species of the Mediterranean Basin. However, given the potential cryptic diversity that may still exist among diverse groups of southern Europe (Praz et al. 2019, 2022), providing barcode references from French vouchers remains highly valuable.

The lack of reference sequences for pollinators poses a serious impediment for biodiversity monitoring that relies on meta- or megabarcoding, as well as for developing non-lethal approaches using environmental DNA (eDNA) detection (Thomsen and Sigsgaard 2019; Makiola et al. 2020). This challenge is further compounded by the need for alternative DNA barcode markers in cases where COI is either insufficiently resolutive or unsuitable for analyzing degraded eDNA (Elbrecht et al. 2016; Marquina et al. 2019; Allen et al. 2023). Addressing this issue, a recent study generated reference data for the alternative 16S marker, covering 148 species collected in Occitania, South-West France (Marquisseau et al. 2025). However, this highlights the substantial progress still required to achieve comprehensive coverage for the 982 French species of wild bees (indeed, A total of 982 wild bee species are known from France, excluding the domesticated species, *Apis mellifera* LINNAEUS, 1758; Ropars et al. 2025).

In the present article, we provide 1477 reference barcodes covering ∼560 French bee species. This contribution has increased coverage from 62% to 82% of French species with barcodes and from 25% to 72% with French voucher specimens since the beginning of the project in 2021. This was made possible through the collaboration with sixteen specialists and institutions who opened their collections for tissue material sampling. Relying on experts’ collections provides access to broader taxonomic coverage and highly reliable identifications within a short timeframe (Kotrba 2020), addressing the need for rapid database completion. Collaboration with experts was also crucial for the subsequent validation of the data and for addressing inconsistencies that arise between morphological identification and molecular specimen identification. Thus, in this study, our objectives were: 1) to provide COI-5P reference barcode data for the French bee fauna, specifically species from the Mediterranean Basin, 2) to confirm the global suitability of the COI-5P marker for bee species delineation, despite some groups not exhibiting a proper barcode gap, 3) to highlight inconsistencies opening avenues for taxonomic investigations in the coming years, and 4) to propose a rigorous procedure for establishing reliable DNA barcode libraries based on collection specimens.

### Construction and content

The reference library herein presented was implemented following the procedures described in Figure 1, that we suggest as good practices for the establishment of reference barcode libraries. Key steps are detailed below, they involve: 1) Assessment of existing data and identification of gaps, 2) Data acquisition through biological material sampling and sequencing, 3) Sequence analyses to confirm reference barcodes.

**Figure 1:**
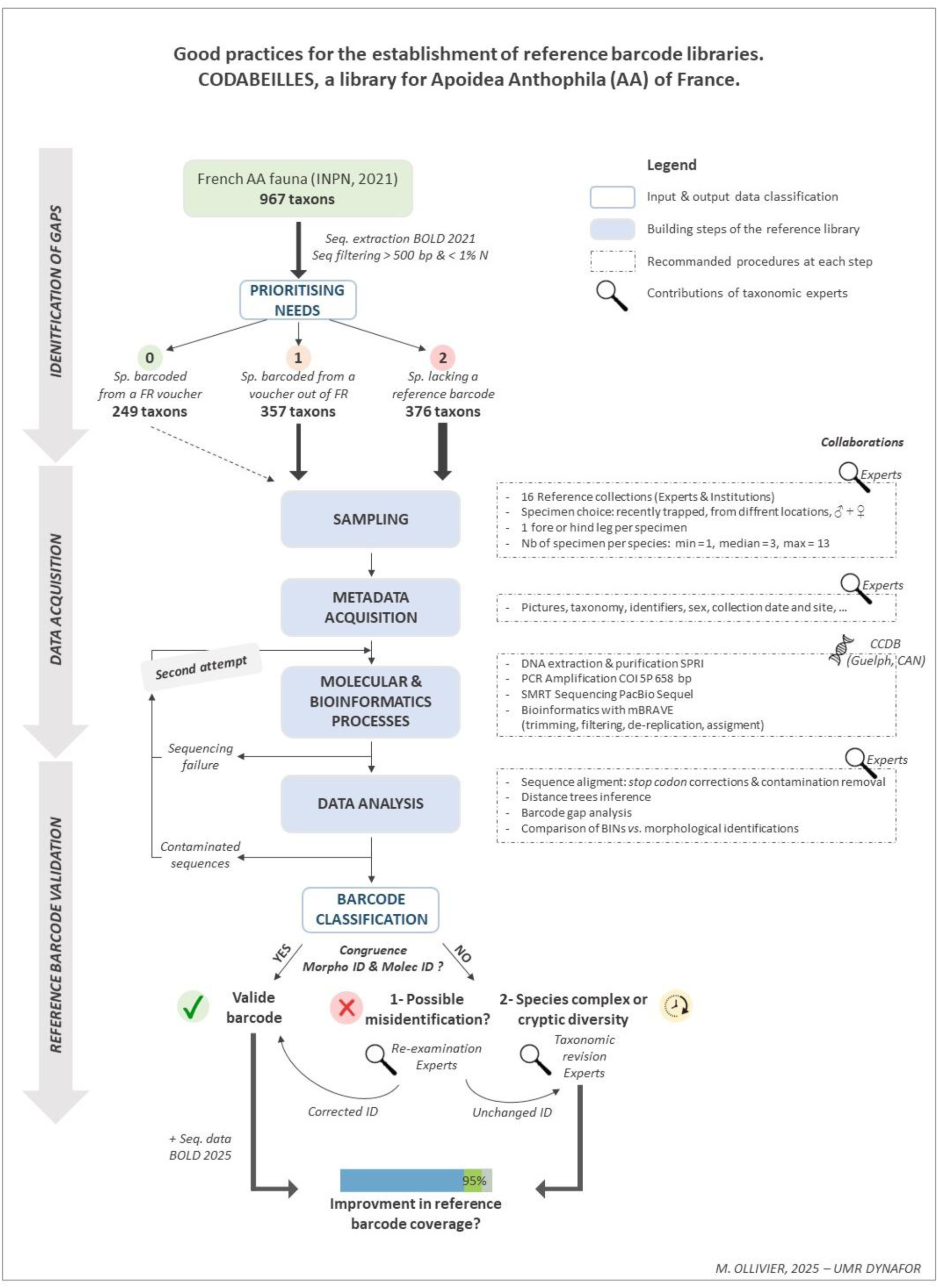
Good practices for the establishment of reference barcode libraries.

#### 1) Assessment of existing data and identification of gaps

- *Existing data extraction:* In spring 2021, the checklist of the wild bee species of France (Hymenoptera: Apoidea: Anthophila) was obtained from the taxonomic repository (TAXREF version 15.0; Gargominy et al. 2021) implemented by the National Inventory of Natural Heritage. The domestic honey bee, *Apis mellifera*, was excluded from the list and for the remaining 967 species, the public database BOLD system was queried to assess the coverage in reference barcode sequences. Sequence data extraction was done using the R package *bold*. Only a subset of sequences meeting barcode standards were kept to assess species sampling priority (sequences > 500 bp-long, < 1% undefined base and country specification).
- *Prioritising needs:* Three priority levels were defined: level 0 for low priority species (species covered by at least one barcode sequence obtained from a French voucher), level 1 for intermediate priority species (species covered by at least one barcode obtain from a voucher outside of France) and level 2 for high priority species (species lacking a reference barcode). The samplings focused on priority level 1 and 2 in order to 1) improve barcode coverage for French wild bee fauna and 2) better characterize intraspecific genetic diversity. We also considered a few low-priority species when collaborators needed molecular validation of a specimen’s identification.

#### 2) Data acquisition: biological material sampling and sequencing

- *Biological material sampling*: Biological material for DNA extraction was exclusively obtained from dry collection specimens, following the procedures recommended in Ollivier et al. (2025). Sixteen contributors from the CODABEILLES Consortium (supplementary material 1), including institutional and personal reference collections, provided tissue material. Targeting the specimens from collections of experts was preferred to the sampling of fresh material. Expert collections often gather male and female specimens of a species, specimens belonging to rare species and specimens sampled through large temporal sequences offering choice for tissue sampling. Beyond the fact that it avoids killing new specimens to find the right species, collaborating with experts allows a wide taxonomic range of species to be covered rapidly with highly reliable identified specimens. To account for intraspecific genetic divergence, three specimens per species coming from different locations were sampled (min = 1, median = 3, max = 13). Figure 2 shows sites where specimens used in this study were collected in France. A minimum amount of tissue, *i.e.* a single fore or hind leg (depending on the need to access diagnostic characters) was sampled for each specimen, allowing for subsequent re-observation of the voucher specimen. Using flame-decontaminated forceps, tissue samples were placed in 96-well plates (with the exception of well H12, which was left empty to serve as a negative control).
- *Metadata acquisition:* Pictures of the voucher specimen were taken and metadata (including taxonomic identification, identifier, sex, and sampling date and locality) were recorded in an Excel spreadsheet (Microsoft Corporation, 2025) for BOLD submission. When a specimen could not be confidently assigned to a species with 100% certainty, we retained a group-level name (*e.g., Bombus gr. terrestris*, *Halictus gr. simplex*) and provided taxonomic notes in the metadata explaining the morphological ambiguities relative to existing species descriptions. Metadata and pictures for all samples were uploaded to BOLD, under a dedicated project, called CODAB (publicly accessible, see section Availability of data and materials).
- *Molecular and bioinformatic processes:* Complete 96-well plates were sent to the Canadian Centre for DNA Barcoding (CCDB, Guelph, Canada) for sequencing of DNA barcodes. Molecular operations were carried out according to standard procedures for processing high-throughput samples (protocols available on https://ccdb.ca/resources/). For PCR amplification of the full-length (658bp) DNA barcode region of the COI-5P gene, two methods were employed depending on the age of the voucher specimen. For recently collected specimens, a primer cocktail combining the universal LCO1490/HCO2198 primer pair (Folmer et al. 1994) with the LepF1/LepR1 primer pair (Hebert et al. 2004) was used. This procedure was carried out on 23 plates (2165 specimens) out of 24. One plate (95 specimens), containing decades-old specimens collected between 1905 and 1984 from Robert Delmas’s reference collection, was processed using the primers sets developed by Prosser et al. (2016) for degraded DNA. This primer configuration allows generating up to 12 overlapping amplicons of the standard COI-5P marker (D’Ercole et al. 2021). PCR fragments, for all samples (2260), were sequenced with single molecule real-time sequencing (SMRT) on the PacBio Sequel platform at the CCDB (Hebert et al. 2018). SMRT sequences were analysed using mBRAVE (Multiplex Barcoding Research and Visualization Environment; www.mbrave.net) with a standard pipeline involving sequence trimming, quality filtering, de-replication, identification, and OTU generation.

#### 3) Sequence analyses to confirm reference barcodes

- *Removal of non-biological sequences*: Following molecular and bioinformatic processing, sequences were made accessible on BOLD under CODAB project where users can obtain information on their quality. Some records were flagged due to 1) the detection of stop codon (45 sequences), and 2) the suspicion of contamination (129 sequences). The contaminated sequences were confirmed with batch ID Engine (using analytical tools of BOLD workbench) and eliminated. Using MEGA11 software (Tamura et al. 2021), sequences flagged for stop codon detection were aligned against sequences from conspecifics, or if not available, from closely related species (same subgenus or genus) in order to point out the possible reading frame shift caused by the presence of indels and leading to the introduction of stop codons. Sequences were edited to correct for these minor errors.
- *Bee sequence analyses*: To estimate the relevance of the COI-5P marker for the delimitation of French wild bee species, we implemented classical methods based on sequence divergence. Intraspecific and interspecific variations were assessed with pairwise distances using the Kimura 2-parameter (K2P) distance model (Kimura 1980) and pairwise deletion method. To provide a graphical representation of species divergence, neighbor-joining (NJ) trees were inferred using sequences produced in this study along with sequences already available on BOLD for the same species or genera. To confirm the existence and extent of the barcode gap in French wild bee species, we plotted the maximum intraspecific distances against the interspecific (nearest neighbour) distances. A histogram was generated to illustrate the distribution of normalized divergence at both species and genus level. These analyses were conducted using sequences longer than 400 bp, with less than 1% unidentified bases and for which species identification was not doubtful. For sequences meeting barcode standards (sequences > 500 bp-long, < 1% undefined base) a Barcode Index Number (BIN) was assigned automatically by BOLD (Ratnasingham and Hebert 2013). BINs are sequence clusters that often correspond to a biological species and help to assess the congruence between molecular clusters and morphological identifications. This combination of approaches was necessary to ensure the reliability of sequences as reference barcodes and to highlight some inconsistencies that led experts to re-identify some of the voucher specimens. Several scenarios were observed and are discussed in the next section: (1) congruence between morphological and molecular species-level identification, and (2) incongruence between morphological and molecular results, which may be due to putative misidentifications of the voucher specimen, specimen belonging to a known species complex, unstable taxonomy, or suspected cryptic diversity. Records attributed to these cases made up the actual CODABEILLES library, a dataset accessible through BOLD at http://dx.doi.org/10.5883/DS-CODAB01.

Eventually, to account for possible barcodes generated outside the CODABEILLES initiative also contributing to French wild bee species coverage over the same period, the BOLD checklists tool was used on the 982 French wild bee species (Ropars et al. 2025).

**Figure 2:**
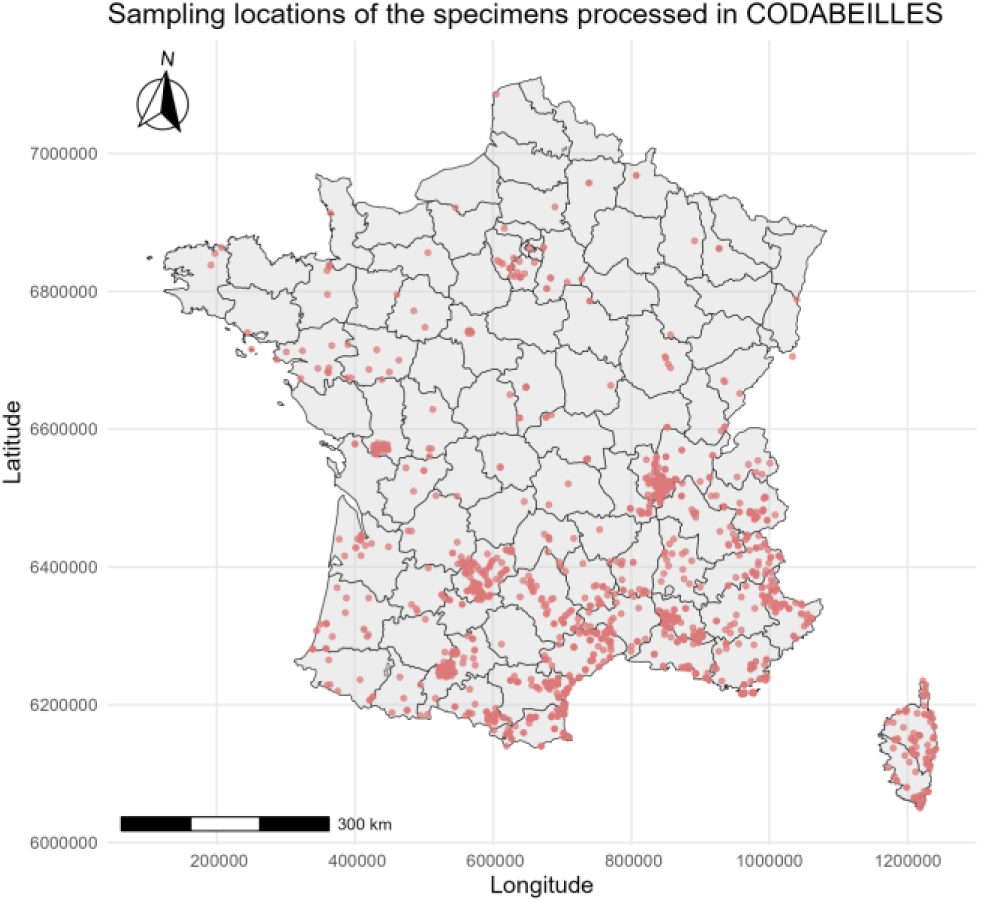
French sampling locations of the specimens processed in CODABEILLES project. 34 specimens were collected in other countries but corresponded to species found in France (4 in Belgium, 3 in Greece, 7 in Italy, 4 in Morocco, 3 in Portugal, 7 in Spain, 3 in Switzerland, 2 in Tunisia).

### Utility and discussion

The reference library, generated under the CODABEILLES initiative for French wild bees, is available on the BOLD Public Data Portal. It can be downloaded in TSV format, including metadata and sequences, at http://dx.doi.org/10.5883/DS-CODAB01. The data follow the standardized format provided by the Barcode Code Data Model Records (see the GitHub page: https://github.com/boldsystems-central/BCDM/tree/main and for details on field descriptions, refer to the *<u>field_definitions.tsv</u>* file). Records are also searchable within BOLD using the search engine. As an alternative way to explore and use these reference data, users can log in and access the library dataset directly through BOLD via the workbench platform.

#### 1) Sequencing success from specimens in collection varies across taxonomic groups

The 2260 specimens provided by the 16 institutional and private collections represented 649 taxa (630 species formally identified morphologically and 19 ambiguous taxa) across 56 genera within the six European bee families. After filtering out contaminations and low-quality sequences, 1477 sequences were retained, belonging to 561 taxa and 53 genera (Table 1). Among the 2165 specimens processed using the standard protocol targeting the 658 bp length COI-5P marker, the overall sequencing success rate reached 70%. A total of 1439 sequences met barcode standards (>500 bp in length, <1% ambiguous bases), while 10 sequences were shorter, ranging from 444 to 499 bp. This result is quite encouraging, given that most collections providing bee tissue samples lacked storage conditions specifically designed for DNA preservation, which may have contributed to amplification or sequencing failures in some samples (Prosser et al. 2016; Nakahama 2021). The sequencing success rate also aligns with previous studies using comparable molecular approaches, which reported similar recovery rates ranging from 67% (Schmidt et al. 2015) to 83% (Villalta et al. 2021), the latter based on specimens sequenced within four years after their capture in the field. However, despite employing a NGS approach suitable for old material (Prosser et al. 2016), only 28 partially informative sequences (ranging from 130 bp to 456 bp) were recovered from the 95 specimens in Robert Delmas’s reference collection (*Andrena* spp. and *Bombus* spp. collected between 1905 and 1984).

**Table 1:** Overall data regarding the CODABEILLES library. Number of bee specimen sampled, Number of sequences generated (>500 bp and <500 bp), Number of taxa and genera sampled and covered with a sequence. Taxa refers to either species or species groups in cases of uncertain species assignment.

|  |  |
| --- | --- |
| Nb of sampled specimens | **2260** |
| Nb of sequences generated | **1477** |
| Nb of sequences meeting barcode standard (>500 bp & <1% N) | **1439** |
| Nb of partial sequences (<500 bp) | **38** |
| Nb of taxa with a sequence/Nb of taxa sampled | **561/649** |
| Nb of genera with a sequence/Nb of genera sampled | **53/56** |

Combining both approaches enabled sequence coverage for 86% of the sampled taxa (Table 1). As reported in previous studies, sequencing success was variable amongst bee taxonomic groups (Magnacca and Brown 2012; Sheffield et al. 2017; Villalta et al. 2021). For instance, in this study, 81% of Megachilidae specimens were successfully sequenced, highlighting that the molecular protocol was particularly effective for the genera *Osmia* and *Megachile*, with success rates of 90% and 87% respectively (Table 2, *b)*). In contrast, only 43% of the specimens belonging to genus *Hylaeus* and 49% belonging to genus *Andrena*, provided a sequence (Table 2, *b)*). In the worst-case scenarios, despite multiple attempts, certain species proved particularly resistant to amplification (Table 2, *c)*). In the case of *Andrena* and *Lasioglossum*, this low success rate may be explained by inefficient primer annealing due to either polymorphic sequences or single nucleotide insertion or deletion within the primer binding site resulting in a lower binding efficiency or non-specific amplifications (Magnacca and Brown 2012). Otherwise, it may be attributed to the presence of heteroplasmy in *Hylaeus* species (Magnacca and Brown 2010). In such cases, alternative primer pairs or barcode markers are recommended. The approach implemented by Wood et al. (2024) for Iberian bees — using two different primer pairs to amplify overlapping fragments of 325 bp and 418 bp within the 658 bp standard COI-5P gene — successfully improved DNA amplification for five of these recalcitrant species (*but see section 4 Planned future development*). Additionally, targeting the complementary 16S marker may help overcome this limitation, as reference barcodes for two of these recalcitrant species were recently deposited in the BOLD public database (Marquisseau et al. 2025)

**Table 2:** Sequencing success rates calculated for a) each taxonomic family, b) the 10 most abundant genera, and c) the 9 most problematic species within the CODABEILLES project. These success rates can range from 0% (no successful sequencing) to 100% (all specimens successfully sequenced) and are visually represented using a color gradient from red (null or low success) through yellow (moderated success) to green (high success).

| a) Sequencing success rate per family |  |  |  |
| --- | --- | --- | --- |
| Family | Nb of sampled specimens | Nb of sequenced specimens | Success rate (%) |
| Andrenidae | 480 | 235 | 49 |
| Apidae | 799 | 552 | 69 |
| Colletidae | 133 | 77 | 58 |
| Halictidae | 402 | 257 | 64 |
| Megachilidae | 401 | 326 | 81 |
| Melittidae | 45 | 30 | 67 |

| b) Sequencing success rate for the 10 most abundant genera in the CODABEILLES project |  |  |  |
| --- | --- | --- | --- |
| Genus (family) | Nb of sampled specimens | Nb of sequenced specimens | Success rate (%) |
| *Andrena* (Andrenidae) | 458 | 224 | 49 |
| *Nomada* (Apidae) | 223 | 180 | 81 |
| *Bombus* (Apidae) | 209 | 127 | 61 |
| *Lasioglossum* (Halictidae) | 184 | 111 | 60 |
| *Osmia* (Megachilidae) | 103 | 93 | 90 |
| *Anthophora* (Apidae) | 97 | 67 | 69 |
| *Megachile* (Megachilidae) | 92 | 80 | 87 |
| *Hylaeus* (Colletidae) | 90 | 39 | 43 |
| *Eucera* (Apidae) | 86 | 53 | 62 |
| *Sphecodes* (Halictidae) | 85 | 49 | 58 |

| c) The 9 most problematic species for sequencing of the standard CO1 marker in the CODABEILLES project |  |  |  |
| --- | --- | --- | --- |
| Species | Nb of sampled specimens | Nb of sequenced specimens | Success rate (%) |
| *Lasioglossum bimaculatum* | 12 | 0 | 0 |
| *Andrena tenuistriata* | 10 | 0 | 0 |
| *Lasioglossum prasinum* | 10 | 0 | 0 |
| *Chelostoma florissomne* | 9 | 0 | 0 |
| *Andrena aeneiventris* | 7 | 0 | 0 |
| *Bombus jonellus* | 7 | 0 | 0 |
| *Andrena strohmei* | 6 | 0 | 0 |
| *Megachile pilicrus* | 5 | 0 | 0 |
| *Andrena agilissima* | 13 | 1 | 8 |

#### 2) A significant improvement in reference barcode coverage for the French bee fauna

The CODABEILLES initiative has played a key role in expanding the reference library for the six European bee families (Figure 3). At the project’s launch in 2021, only 25% of the French bee fauna was covered with a reference barcode obtained from a voucher from France, increasing to 62% when including data from other countries (Schmidt et al. 2015; Villalta et al. 2021). At the date of writing this manuscript, reference data for French species has significantly improved: 72% of the 982 wild bee species from France (Ropars et al. 2025) are now covered by a French reference barcode. When incorporating reference sequences from projects conducted in other geographical areas over the same period (*e.g.* Iberian bee species, Wood et al. 2024), overall coverage reaches 94% of the French bee fauna.

**Figure 3:**
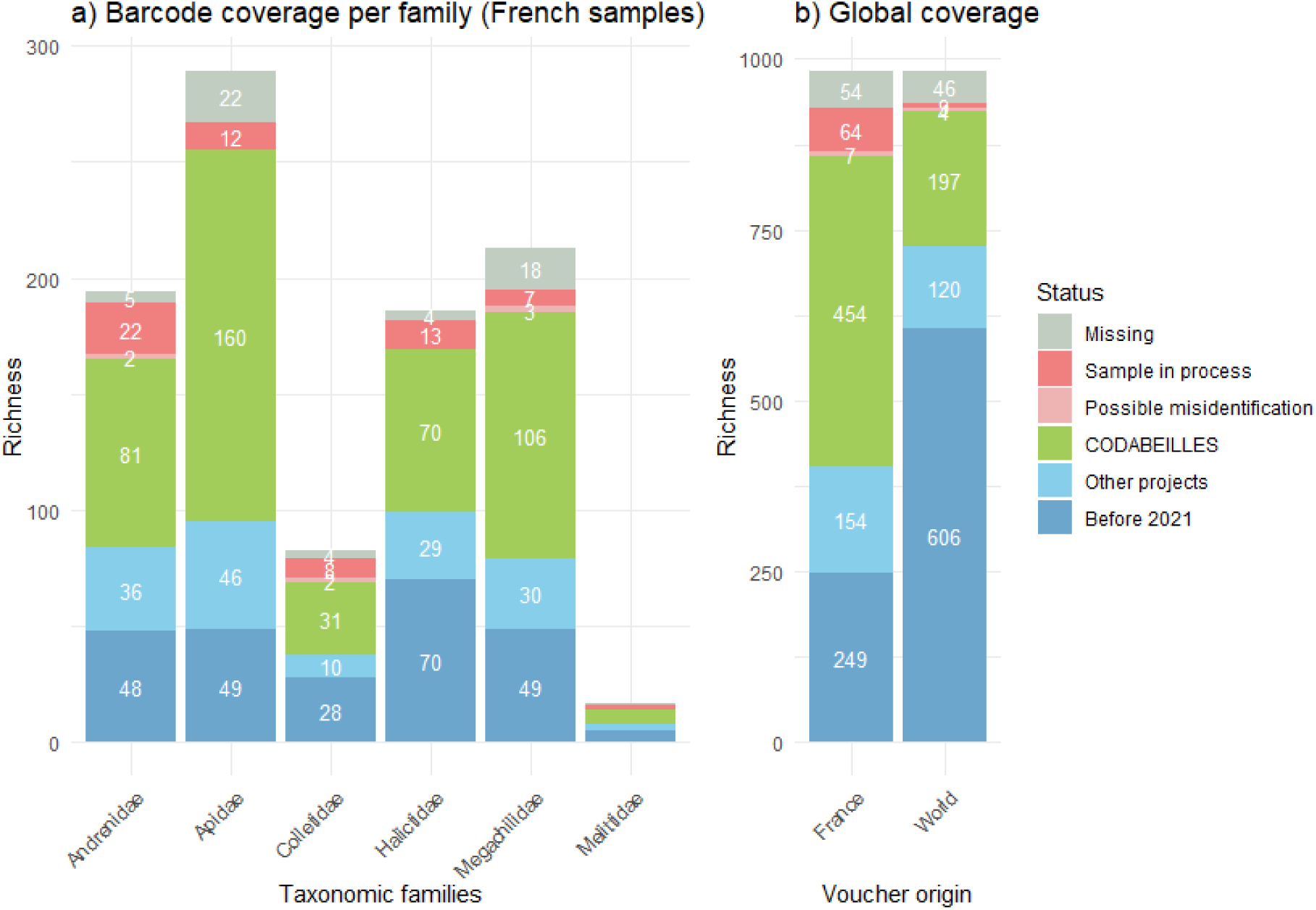
Improvement in barcode coverage for French wild bees since 2021.

Reference sequences were obtained for 561 taxa, including 545 formally identified species (544 French species and one Algerian species). Additionally, 16 taxa could not be accurately identified, as they likely belong to species complexes requiring further taxonomic investigation for unambiguous morphological identification. The supplementary material 2 provides the status of the species as of February 2025, along with the corresponding synonyms under which few specimens have been processed in the CODABEILLES dataset. For four species, the voucher specimens providing reference sequences were collected outside of France: *Amegilla quadrifasciata* (DE VILLERS, 1789) obtained from Morocco; *Andrena subopaca* NYLANDER, 1848 obtained from Belgium; *Nomada fallax* PÉREZ, 1913 obtained from Portugal; and *Nomada numida* LEPELETIER, 1841 obtained from Italy. This information is accessible in supplementary material 2 and 3. The CODABEILLES initiative contributed new French reference barcodes for 454 of the 982 French species (Ropars et al. 2025), 197 of which had no public reference barcode in the BOLD world database (Figure 3). It also provided additional reference sequences that improved the characterization of intraspecific genetic divergence for 78 species. Alongside the recent work of Wood et al. (2024), this project delivers new and complementary reference data to enhance taxonomic knowledge on the bee species from the Mediterranean Basin (Wood et al. 2024; Schneider et al. 2024).

A total of 59 wild bee species (6%) are still lacking barcode reference for the COI-5P marker. This includes species across the six European bee families: eight Andrenidae species, 21 Apidae, eight Colletidae, six Halictidae, 15 Megachilidae, and one species from Melittidae (Figure 3). This number includes four species with a barcode but for which we lack confidence in the identification of the voucher. At a time when nature conservation is a priority, taxonomic research on bees is particularly active in Europe. Revisions are occurring across most genera and families, leading to frequent updates of wild bee checklists (Müller 2020; Dufrêne 2021; Wood et al. 2021; Reverté et al. 2023; Ghisbain et al. 2023; Aubert and Leclercq 2024; Dorchin and Michez 2024; Rasmont and Wood 2024; Aubert et al. 2024; Le Divelec et al. 2024; Le Divelec 2024; Ropars et al. 2025). Of the 59 species lacking a reference barcode, a significant part was either recently discovered on the French territory or recently described as new to science (16, including *Lasioglossum inexpectatum* FLAMINIO & PAULY, 2024; *Aglaoapis sparsepunctata* LE DIVELEC, 2024; *Chelostoma incisa* LE DIVELEC, 2024; *Hoplitis agnielae* LE DIVELEC, 2024; *Hoplitis corsaria* (WARNCKE, 1991); Le Divelec et al. 2024; Flaminio et al. 2024; Le Divelec 2024). They were included late or not included in the CODABEILLE initiative. Other taxa were rare (16 cleptoparasitic bees and notoriously rare taxa such as *Thyreus hellenicus*, Ropars et al. 2025) or we faced sequencing or identification issues with the specimens we sampled (13 cases). We are currently trouble-shooting the sequencing issues and we may obtain new barcodes for up to nine additional species (*Section 4 Planned future developments* and Supplementary material 2).

#### 3) COI DNA barcoding is effective for the delineation of most French bee species but highlights outstanding taxonomic inconsistencies

Out of the 1477 sequences obtained in the CODABEILLES dataset, a total of 1051 sequences exhibited congruence between morphological and molecular classifications, where each BIN was assigned to a single species and each species corresponded to a single BIN (Supplementary material 3). Excluding putative misidentifications and short sequences that could not be assigned to a BIN, this accounted for 76% of the bee samples, enabling the unambiguous classification of 78% of the taxa in the CODABEILLES dataset (*i.e.* 437 species). The scatterplot (Figure 4, *a)*) illustrates the overlap between maximum intraspecific distances (singleton excluded) and interspecific (nearest-neighbour) distances, for 540 pairwise comparisons based on 1386 sequences (length >400 bp). With this subset of data, we found that for 90% of these pairwise comparisons the distance to nearest-neighbour species exceeds the maximum intraspecific distance by at least 2%. Overall, these results suggest that COI DNA barcoding would be a valuable approach for the identification of specimens and the delineation of a majority of bee species (437 species in the present study), as reported in comparable studies that partially covered the French bee fauna (Schmidt et al. 2015; Villalta et al. 2021). However, our results are less conclusive regarding the relevance of COI DNA barcoding compared to findings for Iberian bees, where 95% of specimens identified at the species level were assigned to unique BINs (Wood et al. 2024).

**Figure 4:**
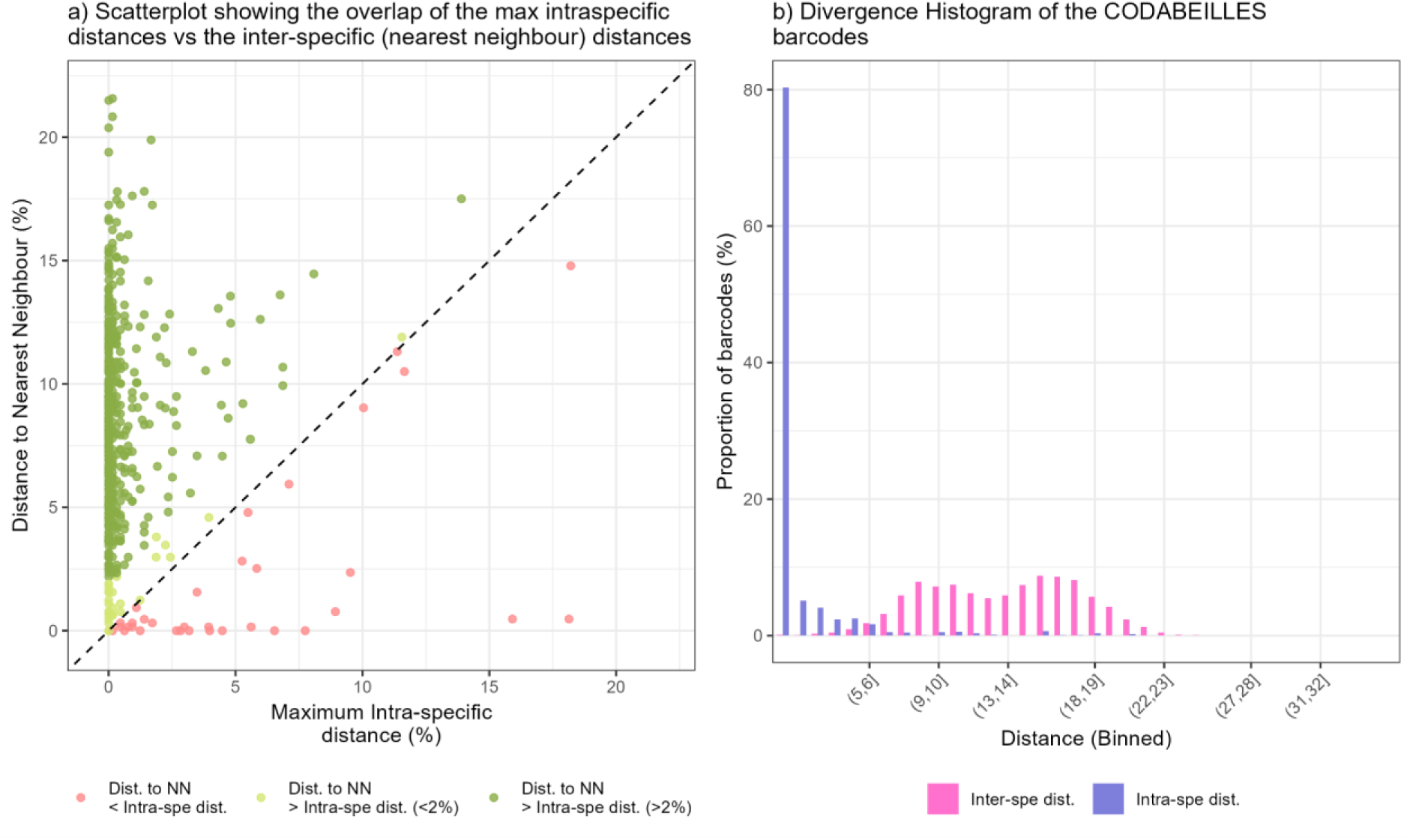
Barcoding gap analysis. a) Scatterplot showing the overlap of the max intraspecific distances against the interspecific (nearest neighbour) distances. b) Nomalized Divergence Histrogram.

Indeed, these conclusions should be interpreted with caution, as we also observed clear cases of taxonomic discrepancies. Some specimens morphologically assigned to a single species were split into multiple BINs, while others shared the same BIN despite belonging to distinct species (Supplementary material 3). Among the mismatches between morphological identifications and molecular results (BIN assignment), 333 were linked to taxonomic inconsistencies probably due to unstable systematics—such as unclear species boundaries within known species complexes or potential cryptic diversity—while 66 mismatches were attributed to potential misidentifications or labelling errors. As of the writing of this manuscript, 145 specimens have been re-examined, resulting in either species reassignment for the specimens (75 specimens) or confirmation of the initial identification (70 specimens), further highlighting unresolved questions regarding species delineation in certain groups. More details on these discrepancies within the genera *Andrena* and *Nomada* are provided below. Phylogenetic relationships remain unresolved for many bee species. Generic and subgeneric concepts have not been consistently revised, and robust phylogenies are lacking for a substantial portion of bee taxa (Engel et al. 2020). However, taxonomic identification keys are generally available for most described species. In unresolved species complexes, identifications are typically based on a combination of morphological characters, ecological traits, phenology, and distribution data. These sources are used collectively to support identifications, though in some cases uncertainty remains.

The histogram (Figure 4, *b)*) depicts the distribution of normalized divergence at both species and genus levels. Sequence divergence ranged from 0 to 18.21 %, with mean distance of 0.94 % within species, and from 0 to 25.78 % with a mean of 13.18 % within genus. Although a bimodal pattern was observed between intra- and interspecific distances, no clear barcoding gap emerged. A barcoding gap occurs when the intraspecific genetic divergence is an order of magnitude smaller than the interspecific genetic divergence within the group of organisms considered (Meier et al. 2008). For example, one might expect the maximum genetic distance within a species not to exceed 3%, while the minimum distance between species (interspecific distance) would be around 7%. In such a case, a barcoding gap is observed between 3% and 7%. In the present study, intra- and interspecific distances overlapped, indicating that this clear separation was not consistently observed. Some species exhibited unexpectedly high intraspecific divergence, and others showed low (<2%) or even null interspecific distances, potentially rendering DNA barcoding ineffective for their delineation.

The concept of a barcoding gap—typically defined as predefined sequence divergence threshold between species—was initially introduced in the literature as a convenient guideline for species identification, not only for Anthophila but for animals in general (Hebert et al. 2003). A recent study analysing all available COI-5P sequences from BOLD for European bee species reported an overlap between intra- and interspecific genetic distances, indicating the absence of an arbitrary barcoding gap across all Anthophila (Janko et al. 2024), although filtering the distance dataset to remove erroneous sequences helped define a barcoding gap ranging between 6.5% (maximum intraspecific distance) and 9% (minimum interspecific distance). The absence of a well-defined barcoding gap was also reported for Irish solitary bees (Magnacca and Brown 2012) and for species from genus *Lasioglossum* in North America (Gibbs 2018), one of the most species-rich bee genera. These studies on wild bees concluded that such a barcoding gap was biologically unlikely.

This may be attributed to several factors: (1) a sequence divergence threshold specific to a given bee genus, probably due to different coalescence times for each lineage (Čandek and Kuntner 2015; Gonçalves et al. 2022; Janko et al. 2024; Marquisseau et al. 2025), (2) uneven barcoding efforts across geographical regions, leading to underrepresentation of certain areas and their diversity (Praz et al. 2022), and (3) erroneous data due to misidentified specimens in public repositories that biases genetic distance estimates (Čandek and Kuntner 2015; Janko et al. 2024). Indeed, overlap between intra- and interspecific distances frequently emerges with increased sampling across the species’ range. Several studies have suggested that the apparent presence of barcoding gaps may be an artifact of insufficient sampling, especially when the full geographic and ecological range of the species was not adequately covered (Meyer et al. 2005; Wiemers and Fiedler 2007).

As a clade that emerged around 125 million years ago and representing over 20,000 species (Cardinal 2018), bees exhibit extreme diversity, which may be incompatible with the existence of a universal COI DNA barcoding gap. The analysis of intraspecific and interspecific distances at the family level and within the most represented genera in the CODABEILLES dataset revealed varying distribution patterns across taxonomic groups, with no clear barcoding gap observed between intra- and interspecific distances (Supplementary material 4). Mean intraspecific divergence was 2.12 % (range 0–18.14) and mean interspecific divergence was 16.94 % (range 0–24.56) for species from *Andrena*, while mean intraspecific divergence was 0.61 % (range 0–4.32) and mean interspecific divergence was 13.12 % (range 2.34–20.43) for species from *Osmia*. Waiting for a more stable consensus on bee classification, sequence trimming—applied by Janko et al. (2024) on a larger dataset—may help filter taxonomically ambiguous cases and clarify the barcoding gap observed at the European level for the genera *Andrena* (10%–13%) and *Osmia* (2.5%–5.5%).

In the present dataset, the genera *Andrena* and *Nomada* exhibited particularly high levels of mismatch between morphospecies and BINs. They were also the genera covered by the higher number of sequences in the present database (Table 2). The following paragraphs aims to highlight the taxonomic discrepancies observed within these two genera, emphasizing the need for further taxonomic investigations in the coming years.

- *Notable inconsistencies within the* Andrena *genus*

Most of the inconsistencies observed within the *Andrena* genus corresponded to single species splitting into multiple BINs, while few cases referred to different species merging into a single BIN or a mix of both situations (Figure 5). Notably, the specimen BCA0624 morphologically identified as *A. assimilis* RADOSZKOWSKI, 1876 (sampled under the synonym *A. gallica* SCHMIEDEKNECHT, 1883 in the CODABEILLLES dataset) and the specimen BCA0829 identified as *A. thoracica* (FABRICIUS, 1775) were both attributed to BOLD BIN #AAE1815. Public information provided on BOLD indicated that this genetic cluster encompassed mostly female specimens belonging to the following species: *A. thoracica*, *A. limata* SMITH, 1853, *A. nitida* (MÜLLER, 1776). As females of *A. assimilis* are morphologically close to those of *A. thoracica* and *A. limata*, differing on the punctuation of the disc of the first tergite (Wood 2023), we cannot exclude a misidentification of the specimen BCA0624 if these diagnostic character are variable within species. However, the boundaries between these three species are unclear, and Wood (2023) has even recently highlighted the existence of three clades formed by different *A. limata* specimens, with no geographic pattern. Given the current state of systematics, the specimen *A. gallica* (BCA0624) would likely belong to *A. limata* clade *#2* (*sensu* Wood 2023). Meanwhile, the specimen identified as *A. thoracica* (BCA0829), although belonging to the same BIN, would be correctly identified being part of a monophyletic group, distinct from *A. limata* clade *#2* (Supplementary material 5). The identity of these specimens remains to be confirmed, as the situation within this species group could not be clarified using COI alone (Wood 2023). This would require further investigations and raises questions about the robustness of the morphological criteria used to distinguish them.

**Figure 5:**
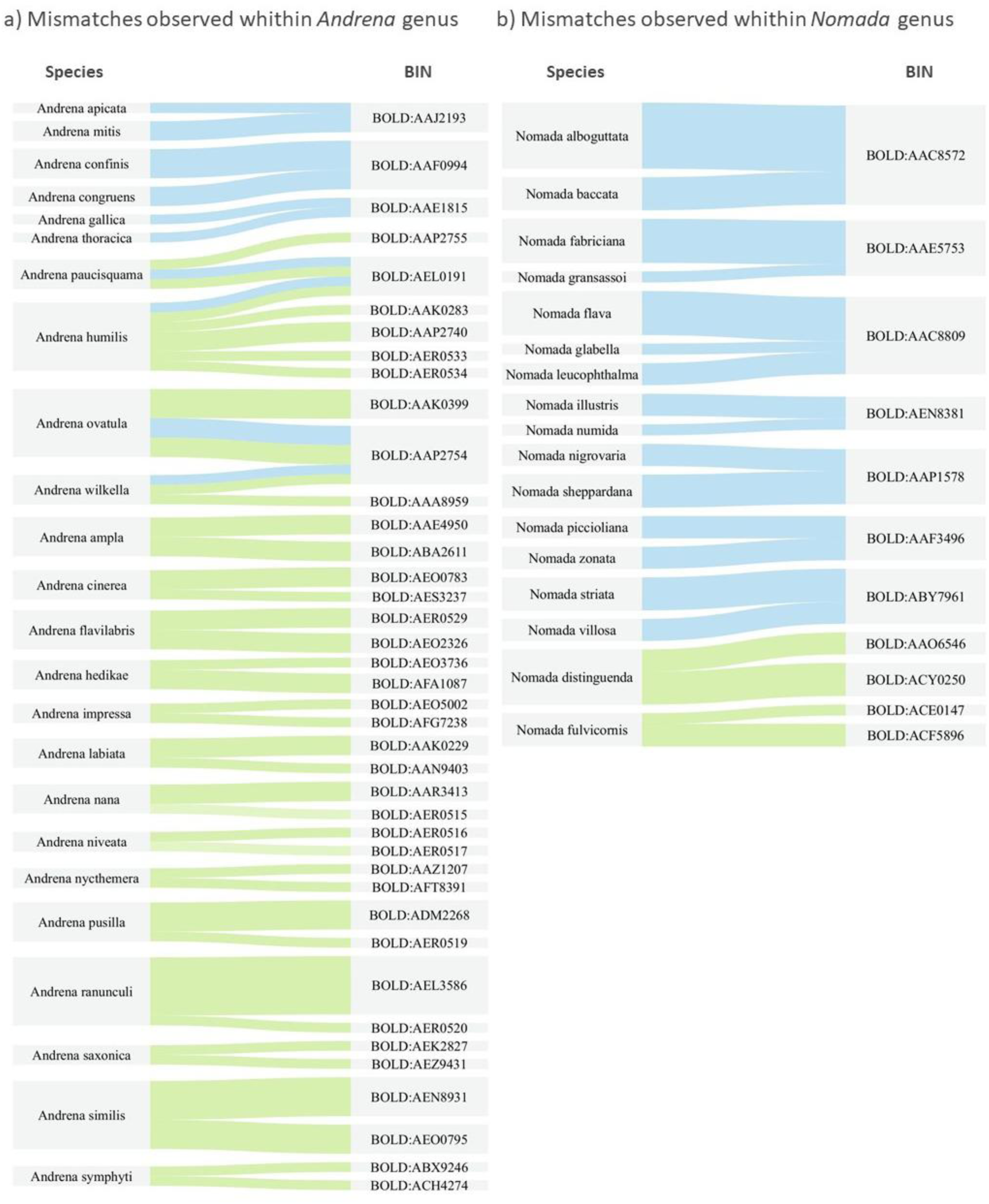
Sankey diagrams illustrating the mismatches observed between morphological and molecular species-level identifications within a) *Andrena* et b) *Nomada* genera. The thickness of the link is proportional to the number of specimens (ranging from 1 to 6). A blue link represents different species sharing a common BIN, while a green link represents a single species splitting into multiple BINs.

Two more pairs of species also shared common genetic clusters: *A. mitis* (NYLANDER, 1852) (2 specimens) and *A. apicata* SMITH, 1847 (1 specimen) under BIN #AAJ2193, as well as *A. confinis* STÖCKHERT, 1930 (3 specimens) and *A. congruens* SCHMIEDEKNECHT, 1882 (2 specimens) under BIN #AAF0994 (Figure 5). Nonetheless, the *A. mitis* specimens exhibited about 0.82% divergence from the *A. apicata* specimen. Despite being grouped under the same BIN, this slight genetic divergence enables their distinction using a reference barcode. This suggests that COI DNA barcoding remains effective for species delineation even below the conventional 2% interspecific divergence threshold, particularly for species that may have undergone a recent speciation event (Bossert et al. 2022). These refer to situations highlighted in light green on the scatterplot (Figure 4, *a)*).

Among the discrepancies observed within the *Andrena* genus, some cases involved mixed patterns (Figure 5). Specimens identified as *A. ovatula* (KIRBY, 1802) and *A. wilkella* (KIRBY, 1802) exhibited both BIN sharing and fragmentation into multiple BINs. This observation was expected, as these species belong to a well-known species complex. Recent taxonomic investigations, using short diagnostic barcodes, have enabled the separation of *A. ovatula* from *A. afzeliella* (KIRBY, 1802), previously identified as *A. ovatula sensu lato* (Praz et al. 2022). The NJ tree inferred using all publicly available sequences from the *Taeniandrena* subgenus helped in clarifying the identity of the specimens from CODABEILLES library (Supplementary material 6). Specimens from *A. wilkella* formed a monophyletic group attributed to BIN #AAA8959. The BIN #AAK0399 corresponded to specimens of *A. ovatula* from diverse origins (Portugal, Spain, UK and France) but exhibiting a very low mean divergence of 0.09%. The BIN #AAP2754 should refer to the *A. afzeliella* genetic cluster, as demonstrated by Praz et al. (2022). This suggests that the following specimens should be re-observed in light of the description of Praz et al. (2022), probably leading to a revision in the species attribution: specimens BCA0343, BCA0225 and BCA0911 (all assigned to BIN #AAP2754) may correspond to *A. afzeliella*.

A second instance of a mixed pattern was observed between specimens of *A. humilis* IMHOFF, 1832 and *A. paucisquama* NOSKIEWICZ, 1924, which yielded a rather unexpected result (Figure 5). One specimen (BCA0743) morphologically attributed to *A. paucisquama* was assigned to BIN #AAP2755 (Supplementary material 7). This genetic cluster consisted of five specimens from various locations (Austria, Croatia, Greece and France) all identified as *A. paucisquama* and showing no genetic variation, an outcome that aligns with expectations (Supplementary material 7). Unexpectedly, however, a second *A. paucisquama* specimen (BCA1634, from Hérault, FR), was assigned to a different BIN (#AEL0191). This BIN also included another specimen (BCA0052, from Gers, FR), that belonged to the species complex composed of *A. humilis* and *A. cinerea* BRULLÉ, 1832. The BIN #AEL0191 exhibited 18.14% divergence from BIN #AAP2755 (*A. paucisquama* cluster), while showing 11.02% mean divergence from BIN #AEO0783 (*A. cinerea* cluster) and 12.19% mean divergence from a clade of five BINs associated with *A. humilis* (Supplementary material 7). Both specimens (BCA1634 and BCA0052) from the BIN #AEL0191 are stored in independent collections and were initially identified by different experts. Given their genetic similarity and the isolated clade they form – distinct from any other genetic reference – they warrant closer examinations, as, if not a case of misidentification, this could be a sign of cryptic diversity. Moreover, we observed a situation of high diversification within *A. humilis* with specimens splitting into five different BINs (Figure 5). Wood (2023) and Schmidt et al. (2015) already reported the particularly high intra-specific variations exhibited by this species, and attributed to its range of distribution, *A. humilis* being the most widespread West Palaearctic *Chlorandrena*. In the present dataset, three specimens identified as *A. humilis* supported this broad species concept, being distributed across BINs that group *A. humilis* specimens from Germany, Austria and Belgium (specimen BCA0821 attributed to BIN #AAK0283, as well as specimens BCA0051 and BCA0367 attributed to BIN #AAP2740). However, two more specimens (BCA0369 in BIN #AER0534 and BCA0801 in BIN #AER0533) were strongly separated from the broad *A. humilis* clade by a mean distance of 15.79%. They appeared genetically relatively close to *A. rhenana* STÖCKHERT, 1930 (with a 4.47% mean divergence), while remaining isolated from any other genetic reference and raising further questions about the potential for cryptic diversity. Complementary genetic markers and thorough morphological examination are needed to precisely determine the status of such specimens.

Numerous additional cases of species splitting into multiple BINs were observed within the *Andrena* genus (Figure 5). Specimens of *A. hedikae* JAEGER, 1934 were attributed to two different BINs (distant from 2.04 %). When considering publicly available data, the species formed three clades. One clade contained specimens from Morocco, while the other two included specimens from various locations (Portugal, Spain, France). Although our data provide a better characterization of intraspecific divergence for this species with specimens collected from South West and East France (Gironde and Drôme), additional sequences - especially from South Eastern Europe- are still needed to further elucidate barcode variation in *A. hedikae*, as highlighted by Wood (2023).

Four specimens originally identified as *A. ampla* WARNCKE, 1967 were attributed to two BINs (BCA0772 and BCA0774 in BIN #AAE4950, BCA0614 and BCA0615 in BIN#ABA2611). The *Andrena proxima*-complex has been the focus of recent investigations based on COI and UCE phylogenetic analyses enabling the separation of the species in this group: *A. ampla*, *A. proxima* (KIRBY, 1802) and *A. alutacea* STÖCKHERT, 1942 (McLaughlin et al. 2023). Beyond a genetic differentiation, the three species also display distinct phenologies, *A. proxima* flying significantly earlier in the season compared to *A. alutacea*, while *A. ampla* exhibits an intermediate flight period (McLaughlin et al. 2023). Although the identification of the specimens BCA0614 and BCA0615 were consistent with publicly available data, those of specimens BCA0772 and BCA0774 raised questions as they were genetically similar to specimens of *A. proxima* (null divergence). Moreover, the phenological data regarding these female specimens, collected on a 23^rd^ May (143^rd^ /144^th^ day of the year), aligned with the flight period of *A. proxima*, suggesting that the specimens should be re-examined and their identification revised.

Phenological shifts among closely related species frequently occur within the genus *Andrena* (McLaughlin et al. 2023). Although *A. flavilabris* SCHENK, 1874 and *A. decipiens* SCHENK, 1861 are nearly indistinguishable, a genetic study confirmed a recent speciation event and the existence of the two taxa, a spring-flying species (*A. flavilabris*) and a summer-flying species (*A. decipiens*) (Mandery et al. 2008). However, the sequences provided in the present study did not align with expectations (Figure 5). The four specimens originally identified as *A. flavilabris* were assigned to distinct BINs that exhibited a 2.56 % mean divergence (BCA0348 and BCA0750 in BIN #AER0529, BCA0349 and BCA0350 in BIN #AEO2326). The BIN #AEO2326 exhibited null genetic variation and was also composed of specimens identified as *A. decipiens* from Italy and Spain (publicly available data). The collection dates for *A. decipiens* specimens ranged from 1^st^ July to 9^th^ July. In contrast, the specimens BCA0349 and BCA0350 were collected on 16^th^ of April and 18^th^ of May, respectively, which is consistent with their assignment to *A. flavilabris*, but is rather unexpected since they share similar barcode sequences with *A. decipiens*. The genetic differentiation observed by Mandery et al. (2008) between the two taxa was carried out using a 500 bp fragment of the 16S rRNA gene. In this case of evolutionary young taxa separation, it is possible that the standard COI barcode lacks diagnostic substitutions, making it unresolutive. Besides, and more surprisingly, the specimens BCA0348 and BCA0750 were the only representatives of the new BIN #AER0529. This unexpected level of intraspecific variation did not appear to be linked to their geographical origin (both collected in Rhône, FR) nor explained by their collection dates (17^th^ October and 20^th^ April, respectively). However, this raises questions about the potential bivoltine nature of the species. Further investigation is needed, involving barcoding of additional individuals to better characterize the extent of this variation.

High levels of intra-specific variations within a nominal species may draw attention to cases of unresolved taxonomy, differentiation of isolated populations or potential cryptic diversity. For instance, in the present data, several cases of newly divergent BINs were observed within species (Figure 5). In *A. lavandulae* PÉREZ, 1902 (sampled under the synonym *A. impressa* WARNCKE, 1967 in the present dataset), specimen BCA1575 formed the new BIN #AFG7238, exhibiting a 2.25% mean divergence from BIN #AEO5002 which included multiple conspecific specimens from Spain, Portugal, Morocco and France. In *A. niveata* FRIESE, 1887, specimen BCA0394 formed the new BIN #AER0517, showing a 2.39% mean divergence from the BIN #AER0516. In *A. ranunculi* SCHMIEDEKNECHT, 1883, specimen BCA0783 formed the new BIN #AER0520, diverging by 5.96% from the BIN #AEL3586, which comprises conspecific specimens from Spain and France (notably, BCA0782 was collected at the exact same location on the same date). In *A. nana* (KIRBY, 1802), specimen BCA0772 formed the new BIN #AER0515, distant by 6.19% from the BIN #AAR3413, which contained conspecific references. Collected in Eastern France (Isère), specimen BCA0772 represented the first genetic record from this geographical area, while specimens from BIN #AAR3413 originated from Germany, Portugal, Morocco, Spain and South France (Gers). In *A. pusilla* PÉREZ, 1903, specimen BCA0758 also raised questions. It formed the new BIN #AER0519, exhibiting a high divergence (11.92%) from conspecific specimens assigned to BIN #ADM2268, yet showing relative genetic proximity (5.13%) to BINs #AAV9726 and #ADZ6919, which included specimens from *A. simontornyella* NOSKIEWICZ, 1939. The morphological re-examination of specimen BCA0758 did not align with the genetic findings and, at this stage, failed to provide further clarification.

These cases may reflect overlooked isolated populations or even possible undescribed species, further supporting the theory of rapid diversification reported in the *Andrena* genus (Praz et al. 2022; Wood 2023). Nonetheless, the classification of specimens from a single species into multiple genetic clusters alone is insufficient to draw conclusions on new species boundaries. It is possible that the highlighted specimens represent additional cryptic species, however proper case-by-case investigations are needed, including the genotyping of particularly divergent populations along with the integration of complementary data (phenology, ecological, distribution) within an integrative taxonomy framework. By providing a more detailed characterization of inter- and intraspecific genetic variations, this study establishes a solid baseline for future taxonomic revisions of French bee species distributed across the Mediterranean Basin.

- *Notable inconsistencies within the* Nomada *genus*

All *Nomada* specimens with inconsistent barcode results (Figure 5) were re-examined by a taxonomist of the group (Eric Dufrêne). Specimen BCA1335, initially identified as *N. mutabilis* MORAWITZ, 1870, was the only specimen invalidated (but see details below). Most of the inconsistencies observed within the *Nomada* genus corresponded to different species merging into a single BIN while few cases referred to single species splitting into multiple BINs (Figure 5).

*Nomada alboguttata* HERRICH-SCHÄFFER, 1839 (6 specimens) and *N. baccata* SMITH, 1844 (3 specimens) were grouped within the same genetic cluster (BIN #AAC8572). However, a slight interspecific divergence was observed, separating the species into two clades with a mean genetic distance of 0.36%. This finding aligns with the results of Mignot (2020), who analyzed specimens from the same *N. baccata* population (collected by E. Dufrêne in Yvelines, France) using both a unilocus mitochondrial COI marker and multilocus nuclear UCEs. *Nomada alboguttata* is known to consist of three forms with phenological lags and different hosts for the first two forms, while the host of the third form remains unknown (Schwarz et al. 1996; Smit 2018). *Nomada baccata* is clearly distinguished from *N. alboguttata* by its later flight period, different host, and subtle but consistent morphological differences. Although the interspecific distance is below 2% (Figure 4 *a*), see light green dots on the scatterplot), COI DNA barcoding can still serve as a diagnostic tool to distinguish between the two species.

A similar situation was observed for *N. zonata* PANZER, 1797 and *N. piccioliana* MAGRETTI, 1883. While these two species are morphologically similar, they can be reliably distinguished by a specialist. Despite sharing the same BIN (BIN #AAF3496), they formed two distinct clades with an average genetic divergence of 0.69% (Supplementary Material 8). This low but consistent divergence allows for their genetic differentiation. A notable point to highlight is that one specimen, BCA2251 collected in Corsica, and belonging to the subspecies *N. zonata pulcherrima* STOECKHERT, 1944 is genetically identical to the nominate subspecies *N. zonata zonata* PANZER, 1797.

Likewise, one specimen (BCA0939) identified as *N. numida manni* MORAWITZ, 1877, from Sardinia (IT) was assigned to the same BIN as specimens of *N. illustris* SCHMIEDEKNECHT, 1882, from France, Spain, and Portugal. The two species were separated into two clades by a mean distance of 1.08%, and exhibited clear morphological differences, despite belonging to the same species group (*sensu* Alexander 1994; Alexander and Schwarz 1994) and the same subgenus (*sensu* Straka et al. 2024).

We also observed that two specimens of *N. villosa* THOMSON, 1870 were grouped in the same BIN as *N. striata* FABRICIUS, 1793, along with other publicly available specimens from both species. These two species are genetically close (Straka et al. 2024) and also morphologically close, forming a small, highly homogeneous group alongside *N. symphyti* STOECKHERT, 1930. Interestingly, in this BIN (#ABY7961), we observed three distinct clades (Supplementary Material 8): (1) the two *N. striata* specimens from Corsica, which also differ in their coloration; (2) the *N. striata* specimen from the Pyrénées-Orientales, grouped with Iberian specimens; and (3) the remaining specimens, comprising a mix of *N. striata* and *N. villosa*, from France and other parts of Europe, forming the third clade. This slight genetic differentiation, seemingly linked to geographical patterns and partially supported by morphological variations, would warrant further investigation.

In contrast to previous observations, we found a few cases where a single species was split into two distinct BINs. One notable example is *N. fulvicornis* FABRICIUS, 1793, which was divided into two highly divergent BINs (5.63% genetic distance apart). In BIN #ACE0147, we observed other publicly available specimens of *N. fulvicornis* alongside *N. subcornuta* (KIRBY, 1802), despite the latter’s recent reinstatement as a distinct species—though based on a geographically restricted sampling (Falk 2017; Straka et al. 2024). In the second BIN (#ACF5896), we observed specimens BCA1319 and BCA1320 (collected from Ariège and Dordogne, France, respectively), along with additional *N. fulvicornis* specimens from other projects, originating from Switzerland and Slovakia. Furthermore, two additional genetically close BINs containing *N. fulvicornis* were found in publicly available data. Several subspecies of *N. fulvicornis* have been described, typically with clear geographic differentiation (Falk 2017), but this pattern does not seem to apply in this case. Additionally, multiple morphological forms have been documented, and while some authors consider certain forms to represent valid species, a dedicated study would be required to clarify the taxonomic status of these lineages.

The last notable point concerns specimen BCA1335, initially identified as *N. mutabilis* MORAWITZ, 1872. This specimen formed a new and distinct BIN (#AFA1844) on its own. It is a male belonging to the *N. armata* group (*sensu* Alexander 1994; Alexander and Schwarz 1994), within the subgenus *Gestamen* (*sensu* Straka et al. 2024). All closely related BINs in the cladogram corresponded to species from this group (Supplementary Material 8), which is geographically restricted to the western Palearctic. A detailed morphological examination of BCA1335, in comparison with all known species of the *N. armata* group, suggested that it may represent a new, yet undescribed species (Dufrêne & Philippe, in progress).

#### 4) Planned future developments

Next steps of the CODABEILLES project will involve: 1) the second attempt of amplification for failing samples, 2) the addition of barcode sequences for rare species currently lacking reference data, 3) the re-examination of specimens potentially misidentified or presenting a mismatch between morphological and genetic results and 4) the combination of multiple species delimitation methods to address complex situations.

- Samples that fail to amplify at first attempt targeting the full-length (658 bp) COI-5P gene (716 specimens), will be alternatively processed using internal primers targeting shorter fragments. This approach is recommended in the second instance for degraded DNA of specimens stored in collection (Levesque-Beaudin et al. 2023). Two overlapping fragments of the COI-5P gene, 307 bp and 407 bp-long, will be obtained using two primer cocktail sets: C_LepFolF+MLepR2 and MLepF1+C_LepFolR, respectively (Hajibabaei et al. 2006; Hebert et al. 2013). Once obtained, the new sequences will be deposited under a complementary dataset (DS-CODAB02) on BOLD. This will improve coverage for 64 species with a voucher from France and 9 species with a voucher from another country (Figure 3).
- In the coming months, a focus will be placed on the 59 species currently lacking barcode reference from the French bee fauna. This will be investigated as part of the ongoing related project “IDMYBEES” (https://www.idmybee.com/the-project.html). Since the launch of the CODABEILLES project, close collaboration with experts has been a *sine qua non* for establishing the library, and it will remain essential for achieving this objective. If the target species are not available in existing collections, dedicated field sampling sessions may be required, focusing on specific habitats, flight periods, or host plants. Additional reference barcodes produced will also meet the expectations of international scientific community working on European wild bee species, particularly within the frame of the ORBIT initiative, which aims to develop resources for bee inventory and taxonomy (https://orbitproject.wordpress.com/about-the-project/).
- As discussed above, some specimens will require closer re-examinations, either to revise their identification or to resolve more complex cases of taxonomic discrepancies between morphological and genetic species-level identifications. Such cases of taxonomic discrepancies may result from unclear boundaries within species complexes or a cryptic diversity. In this context, DNA barcodes can serve as a valuable diagnostic character for primary species hypotheses, a first step in a longer integrative taxonomy process (Miralles et al. 2024). For instance, several cases of unexpected genetic variations, confirmed by morphologically distinct characters, were observed within the *Nomada* and *Bombus* genera and should be further investigated. Confirming the existence of a new species will require dedicated studies on additional specimens from these divergent populations, and integrating phenotypic, genetic and ecological data. The use of complementary mitochondrial markers (e.g. 16S; Marquisseau et al. 2025) and nuclear markers (e.g. UCEs; Gueuning et al. 2020) along with the recent sequencing of chromosome-length genomes (Jones et al. 2023; Falk and Mulley 2023; Falk et al. 2025), will enhance resolution and improve the characterization of wild bee diversity. Close collaboration between traditional taxonomy and innovative molecular tools will undoubtedly advance our understanding of wild bee systematics. An integrative taxonomic approach—particularly one that includes phylogenetic inference—would help resolve these species complexes and may reveal new diagnostic morphological features, as demonstrated in the case of the *Lasioglossum villosum* species complex (KIRBY, 1802) (Pauly et al. 2019). This integration is essential for generating reliable reference data, as erroneous information could compromise the effectiveness of future DNA-based monitoring efforts (Locatelli et al. 2020; Janko et al. 2024).
- As recommended within an integrative taxonomy framework, species delimitation should not rely solely on multiple markers but also on the combination of several analytical methods, as each tool presents inherent limitations. In the present study, hypothetical species were grouped based on sequence data using the clustering tool provided by the BOLD platform—specifically, the BIN system (Ratnasingham and Hebert 2013)—on which the barcode data were deposited. However, numerous approaches exist to assess how DNA barcode data from distinct genetic clusters may correspond to biological species. Addressing species complex situations properly would require the integration of dedicated analytical methods within a more comprehensive framework (Lin et al. 2015; Gibbs 2018; Ranasinghe et al. 2022; Vuataz et al. 2024). For instance, the well-known Automatic Barcode Gap Discovery (ABGD, Puillandre et al. 2012) and the tree-based multi-rate Poisson Tree Processes (mPTP, Kapli et al. 2017) are effective methods for delineating species from single-locus data. More recently, the program ASAP was introduced (Puillandre et al. 2021), offering the advantage of providing a score to identify the best species partition. These tools would be particularly well-suited for application to a dataset such as the one presented here.

#### 5) Towards the DNA-based monitoring of wild bee species and their interactions with flowers

The reference data provided by this study serves as a solid foundation for further taxonomic investigations and also constitutes reliable data for the implementation of DNA-based field monitoring of pollinators and their interactions, which is sought at both national and European scales. The incorporation of DNA barcoding, metabarcoding and environmental DNA (eDNA) approaches in biodiversity monitoring offers an opportunity to establish large-scale and long-term monitoring schemes (Piper et al. 2019; Chua et al. 2023). For instance, Creedy et al. (2020) laid the groundwork for using DNA metabarcoding techniques to assess species diversity and abundance. Their study focused on UK bees collected from mass-trapped catches. Steinke et al. (2022) showed that metabarcoding allows for large-scale monitoring of changes in species composition. This approach goes beyond the biomass measurements that have previously been the primary metric for tracking changes in arthropod communities. Moreover, new proof-of-concept methods for non-invasive insect DNA collection have the potential to transform insect monitoring, although further research is needed to assess their scalability and feasibility for routine use. Airborne eDNA metabarcoding demonstrated the ability to detect traces of various pollinators, such as butterflies and wild bees (Roger et al. 2022), while eDNA from flowers has been reported in several studies as capable of revealing a diversity of wild bee species (Thomsen and Sigsgaard 2019; Newton et al. 2023; Avalos et al. 2024). Such an approach would enhance our understanding of species interactions and the ecological and evolutionary processes they support in ecosystems.

## Conclusions

Thanks to the collaboration of sixteen specialists and institutions who provided access to their collections for tissue sampling, we present the CODABEILLES dataset (DS-CODAB01), which includes 1,477 reference barcodes covering approximately 560 French bee species. This contribution has significantly improved barcode coverage, increasing from 62% to 82% of French species with barcodes and from 25% to 72% with French voucher specimens since the project’s launch in 2021. The inclusion of data from other independent initiatives raises the coverage to 94% of the French bee fauna with a barcode, and to 86% when considering only barcodes linked to a French voucher specimen.

Our work allows the identification of most French bee species using their COI-5P gene barcode, paving the way for DNA-based routine monitoring of pollinators. Additionally, our study confirms that Apoidea Anthophila is a highly diverse and taxonomically challenging group, in which species diversity is likely still underestimated. Integrating molecular tools within an integrative taxonomic framework provides experts with new opportunities to contribute to species discovery and classification.

## List of abbreviations

*Not applicable*

## Declarations

### Ethics approval and consent to participate

*Not applicable*

### Consent for publication

*Not applicable*

## Authors’ contributions

MO, AP, and MP conceived and planned the study and coordinated sample collection with partners from the CODABEILLES Consortium. The CODABEILLES Consortium assisted with bee tissue sampling or specimen re-examination. RRo acted as the liaison with the Canadian Center for DNA Barcoding (Guelph) platform. MO, RRo, AP, and MP oversaw data analysis and interpretation. MO and AP analysed the sequences. ED and RRu contributed their taxonomic expertise on the genera *Nomada* and *Andrena*, respectively, and assisted with result interpretation. MO led the manuscript writing by providing the first draft, while ED contributed the initial draft for the *Nomada* section. MO, AM and AP generated figures and tables for the manuscript. All authors reviewed, provided critical feedback, and approved the final manuscript.

The CODABEILLES Consortium is composed of (presented in alphabetical order): Emilie Andrieu^1^, Emmanuelle Artige^2^, Matthieu Aubert^3^, Yvan Brugerolles^4^, Alexandre Cornuel-Willermoz^5^, Raphaël Da Silva Ropio^1^, Adeline Dumet^6^, David Genoud^7^, Benoît Geslin^8^, Laurent Guilbaud^9^, Bernard Kaufmann^6^, Lara Konecny^6^, Anne-Laure Jacquemart^10^, Danny Lebreton^4^, Vincent Leclercq^11^, Gabriel Nève^12^, Annie Ouin ^1^, Christophe Philippe^13^, Bertrand Schatz^14^, Jean-Claude Streito^2^, et Héloïse Vallod^1^.

^1^UMR Dynafor, INRAe, INP-AgroToulouse, Toulouse, France

^2^UMR CBGP, INRAE, CIRAD, IRD, Montpellier SupAgro, Univ Montpellier, Montpellier, France

^3^4 chem. de la Foux, hameau du Méjanel, 34380 Pégairolles-de-Buèges, France

^4^Arthropologia - 60 chemin du Jacquemet, 69890 La Tour-de-Salvagny, France

^5^Office de l’Environnement de la Corse, Observatoire Conservatoire des Invétérés de Corse. 14, Avenue Jean Nicoli, 20250 Corte, France

^6^Université Claude Bernard Lyon 1, LEHNA UMR 5023, CNRS, ENTPE, F-69622, Villeurbanne, France

^7^9 rue Hector Berlioz 87240 Ambazac, France

^8^Université de Rennes (UNIR), UMR 6553 ECOBIO, CNRS, 263 avenue du Général Leclerc, 35042 Rennes cedex, France

^9^UR 406 Abeilles et Environnement Site Agroparc, Domaine Saint-Paul 84914 Avignon Cedex 9, France

^10^Earth and Life Institute, UClouvain, Louvain-la-Neuve, Belgium

^11^5, rue de l’Esplanade, Résidence de l’Arche, Bâtiment Prokofiev, Étage 4, 13090 Aix-en-Provence, France

^12^IMBE, CNRS, IRD, Avignon University, Aix Marseille University, France

^13^Amateur entomologist; Observatoire des Abeilles; 15, rue de l’Auxerrois 46000 Cahors, France

^14^CEFE, CNRS, Univ Montpellier, EPHE, IRD, Montpellier, France

## Acknowledgements

The authors warmly thank Kamila Canale-Tabet, Thibault Leroy, and their research team, as well as Jérôme Willm and Catherine Bonnet, for their valuable assistance with tissue sampling.

## Supplementary materials captions

**Supplementary material 1:**
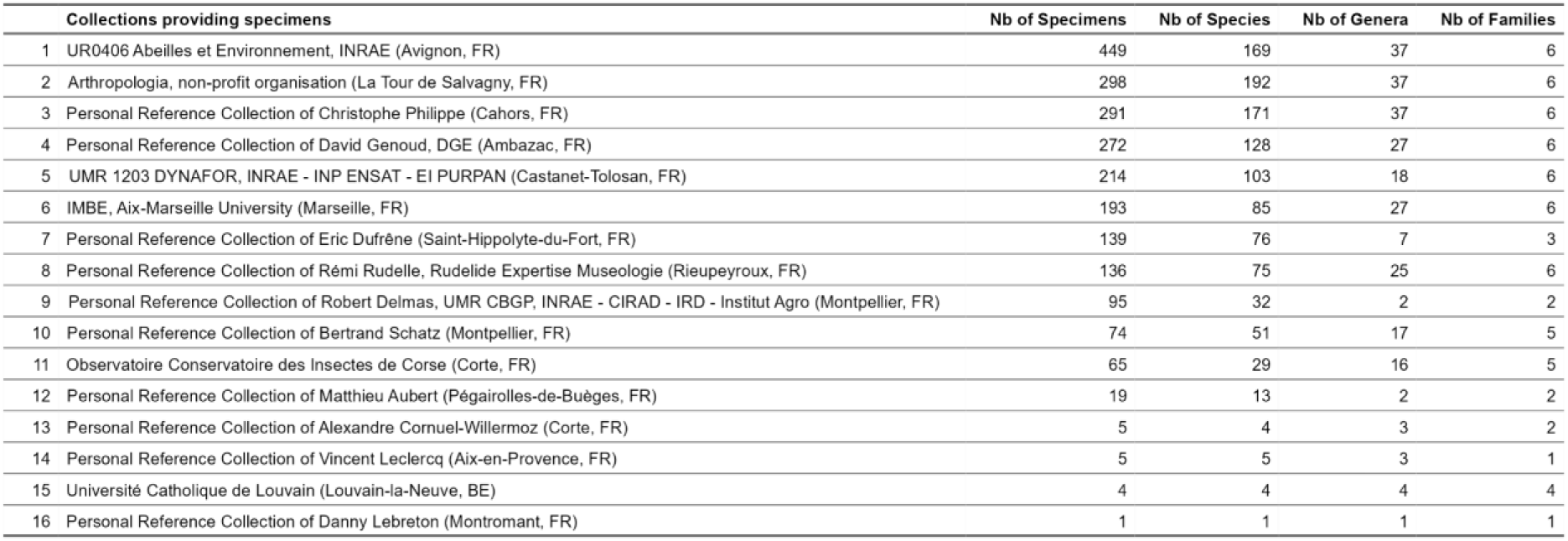
Institutional and personal reference collections providing tissue materials.

Supplementary material 2: Dataset DS-CODAB01

Supplementary material 3: Barcode coverage for French wild bees

**Supplementary material 4:**
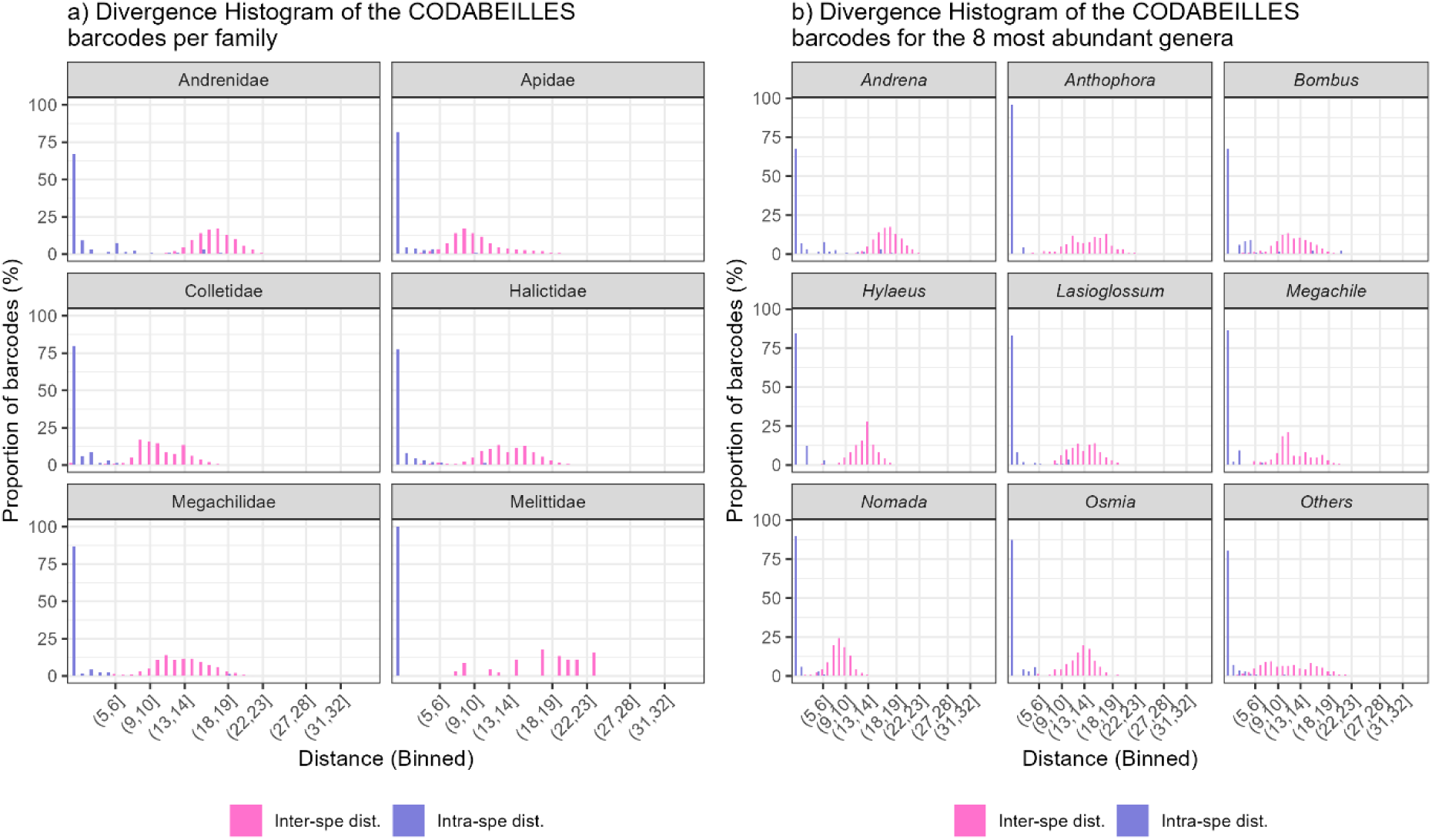
Divergence histograms per taxonomic family and for the 8 most abundant genera.

Supplementary material 5: *Melandrena* subgenus NJ tree including CODABEILLES data along with publicly available barcodes from BOLD.

Supplementary material 6: *Taeniandrena* subgenus NJ tree including CODABEILLES data along with publicly available barcodes from BOLD.

Supplementary material 7: *Chlorandrena* subgenus (including *A. paucisquama*) NJ tree including CODABEILLES data along with publicly available barcodes from BOLD.

Supplementary material 8: *Nomada* genus NJ tree including CODABEILLES data along with publicly available barcodes from BOLD.

